# Putative Cyclospora cayetanensis detection in wastewater metagenomic datasets

**DOI:** 10.64898/2026.09.18.752766

**Authors:** Nicholas R. Minor, William K. Gardner, Marc C. Johnson, Shelby L. O’Connor, David H. O’Connor

## Abstract

Cyclosporiasis is a foodborne illness caused by the parasite *Cyclospora cayetanensis*. Clinical surveillance often lags infection by weeks because diagnosis depends on a clinician ordering a test that is not part of routine parasite panels. Wastewater metagenomic sequencing does not depend on care seeking, which may be useful for early detection and community monitoring. We designed 1,670 31-mers from *C. cayetanensis* mature ribosomal RNA and retained 1,464 without an exact non-target match in the 28 July 2025 NCBI `core_nt` release. By analyzing 293 public wastewater metagenomes, we selected a threshold of 24 distinct diagnostic 31-mers on one read. Applying that threshold to 2,333 public runs from BioProject PRJNA1247874 yielded putative *C. cayetanensis* reads in 81 runs, or 3.5%, and in 24 of the 30 longitudinally sampled sewersheds. Detections rose through the summer in two consecutive years, matching the known seasonality of cyclosporiasis. The most parsimonious explanation is that these reads come from parasites shed into wastewater, although undescribed *Cyclospora* species absent from reference databases cannot be excluded. The same approach can be applied to other pathogens in existing wastewater sequencing data.

## 1 Introduction

Cyclosporiasis is an intestinal illness caused by infection with *Cyclospora cayetanensis*, a single-celled coccidian parasite with no known host other than humans (Casillas et al., 2019). It is acquired by swallowing food or water contaminated with feces. Direct spread from one person to another is unlikely because the oocysts shed in stool are not infectious when they leave the body. Oocysts mature in the environment, a process thought to require at least one to two weeks under favorable conditions. Oocysts are hardy and difficult to destroy. Treating food or water with routine chemical disinfection or sanitizing is unlikely to kill the parasite (Casillas et al., 2019). After ingestion, the parasite settles inside the lining of the small intestine (Ambrose et al., 2025). Many infections produce no symptoms and silent infection is thought to be common in regions where the parasite is endemic. In people who do fall ill, symptoms appear about a week after exposure, with a range of 2 to more than 14 days. Watery diarrhea is frequently accompanied by loss of appetite, weight loss, abdominal cramping, bloating, nausea, fatigue, vomiting, and low-grade fever (Casillas et al., 2019). Left untreated, the illness can persist for weeks or months in a remitting and relapsing pattern before resolving on its own. Trimethoprim-sulfamethoxazole is the recommended treatment (Casillas et al., 2019). Cyclosporiasis can become chronic and cause nutrient malabsorption and weight loss in people who are immunocompromised (Ambrose et al., 2025 and Banasik et al., 2025). Recent genetic analyses have proposed subdividing *Cyclospora cayetanensis* into at least three species (*Cyclospora cayetanensis, Cyclospora ashfordi*, and *Cyclospora henanensis*) (Barratt et al., 2023).

*Cyclospora cayetanensis* was named in 1994 (Ortega et al., 1994), though it was originally described in 1979 (Ashford, 1979) and assigned to the coccidian genus *Cyclospora* in 1993 (Ortega et al., 1993). It emerged as a foodborne threat in the United States during the mid-1990s, and after large multistate outbreaks in 1996 and 1997 cyclosporiasis became a nationally notifiable disease in January 1999 (Casillas et al., 2019). From 2011 to 2015, 2,207 cases were reported, of which 1,384 (62.7%) were acquired within the country and 415 (18.8%) followed international travel (Casillas et al., 2019). Eight states accounted for 1,188 (85.8%) of the domestically acquired cases, with Texas alone reporting 684. All ten outbreaks investigated during that period had illness onsets between May and August, with most symptoms appearing in June or July (Casillas et al., 2019). Travel-associated cases also peak in the same summer months (Casillas et al., 2019).

*Cyclospora* is thought to be endemic in many tropical and semi-tropical regions. Within the United States, contaminated fresh produce drives transmission. Many different products have been implicated including imported basil, raspberries, and snow peas, along with a berry salad, and a prepackaged salad mix. Cilantro may be especially vulnerable to contamination since it was implicated in at least three of the ten outbreaks investigated between 2011 and 2015 (Casillas et al., 2019). Green onions were the likely source of a 2017 cluster tied to a restaurant chain in the Houston metropolitan area (Hall et al., 2020). Cilantro surfaced again in 2023 at a restaurant in Limestone County, Alabama, where 47 people fell ill and illness tracked strongly with cilantro consumption (Goetzman et al., 2025). Since 2018, outbreaks have been traced to produce grown inside the United States (McCaughan and Kniel, 2026). Genotyping by CDC of isolates collected from 2018 to 2021 suggested that there are repeated introductions of the parasite into the food supply rather than an endemic parasite that has settled into a domestic reservoir (Shen et al., 2025).

Detecting *C. cayetanensis* in wastewater has seldom been attempted, and the few studies that have done so rely on a targeted, single-marker approach that identifies the parasite either by oocyst morphology or by amplifying one molecular locus. The first report examined wastewater for the parasite, detecting oocysts microscopically and confirming their identity by molecular techniques (Sturbaum et al., 1998). Later work relied on qPCR applied to bulk sample DNA. A year-long survey of two Arizona treatment plants concentrated water by electronegative filtration and screened the extract with a newly developed quantitative PCR assay (Kitajima et al., 2014). A two-year study of a southeastern Georgia growing region applied the FDA’s validated 18S rRNA gene qPCR (Durigan et al., 2020, published as BAM Chapter 19c) to total DNA extracted from municipal wastewater sludge (Kahler et al., 2024).

This 18S qPCR assay, however, lacks specificity in environmental samples. In the Georgia field study, 26 municipal wastewater sludge samples were flagged by the qPCR assay but only 7 (9% of the 76 sludge samples tested) were confirmed as *C. cayetanensis*, where confirmation required a separate multi-marker genotyping panel of eight nuclear and mitochondrial targets, deep-sequenced and matched against clinical reference haplotypes rather than reading back the short diagnostic amplicon (Kahler et al., 2024). The problem was especially stark in irrigation water: of 65 pond samples flagged by the 18S qPCR, none yielded sequences that clustered with *C. cayetanensis*, the reads instead matching coccidia shed by birds, fish, reptiles, amphibians, and rodents (Hofstetter et al., 2024).

Motivated by the multistate outbreak underway as of July 2026 (Brown, 2026), we reasoned that *C. cayetanensis* could instead be detected directly in existing wastewater metagenomic datasets by requiring exact matches to *C. cayetanensis*-specific 31-mers, drawn from across all rRNA loci.

## 2 Results

Software versions, pinned input identities, and the commands needed to repeat each step on a single machine are collected under REPRODUCING.md.

### 2.1 Ribosomal RNA is a favorable target in PEG-precipitated wastewater

The [CASPER collaborative](https://securebio.org/casper/) collects and performs ultradeep metagenomic sequencing on weekly wastewater samples from sites around the United States (Justen et al., 2026). After filtration to remove solids, particles are concentrated from raw wastewater by polyethylene glycol precipitation, then extracted and sequenced (Rushford et al., 2025). The step was designed to recover virus particles, but polyethylene glycol precipitation is not selective for virions and also brings down particles of similar size and density, including ribosomes. Ribosomal RNA is also recovered, which we have previously leveraged to determine the [vertebrate species contributions](https://dholab.github.io/public_viz/005-12s-species/) to wastewater datasets.

### 2.2 A subtractive design yields 1,670 ribosomal RNA 31-mers absent from all other rRNA

We wanted to identify sequences 31 nucleotides in length (i.e. 31-mers), that occur in *C. cayetanensis* ribosomal RNA and in no other ribosomal RNA. Ribosomes carry mature rRNA, without the spacers and flanking sequence that surround the genes in the rDNA repeat, so mature rRNA sequences are the target for this analysis.

No authoritative set of mature *C. cayetanensis* rRNA sequences exists to use directly. Of the nine reference *C. cayetanensis* rRNA records we could find (config/target_queries.tsv), only AF111183.1 is a complete 18S gene record. The 28S query, MPGL01000046.1, is a 3,519 bp whole-genome shotgun contig from strain CDC:HCRI01:97 rather than a transcript, and the seven `XR_` records for 5S and 5.8S are computational predictions from the RefSeq annotation.

We therefore used these rRNA sequences as the basis for identifying putative mature rRNA sequences queries. Each was aligned with [blastn -task megablast](https://github.com/dholab/cyclospora-in-wastewater-metagenomics/blob/main/01-identify-cyclospora-specific-kmers/scripts/build_rrna_bait.sh#L317-L321) to two pinned *C. cayetanensis* RefSeq assemblies, GCF_002999335.1 (CcayRef3) and GCF_000769155.1 (ASM76915v2), keeping hits at 98% identity and 90% query coverage, and the matching genomic intervals were taken as the target sequence (src/rrna_bait/targets.py). The alignment marks where mature rRNA sits inside the repeat, so spacers and flanks are excluded, and the coordinates of every locus are recorded (curated/target_loci.tsv). The search returned 83 alignment rows at 18 distinct positions in the two assemblies. These were deduplicated to retain two 18S, one 28S, one 5.8S, and three 5S mature rRNA transcripts curated/target_rrna.fasta.

Every 31-mer in those seven sequences and their reverse complements was then computed with [meryl](https://github.com/marbl/meryl), as [meryl count k=31](https://github.com/dholab/cyclospora-in-wastewater-metagenomics/blob/main/01-identify-cyclospora-specific-kmers/scripts/build_rrna_bait.sh#L406-L407). Many 31-mers were duplicated because the two 18S copies are nearly identical and the three 5S copies differ only slightly, leaving 5,561 distinct 31-mers: 1,824 from 18S, 3,457 from 28S, 154 from 5S, and 126 from 5.8S. Every position at which a 31-mer occurs is recorded in a [manifest](https://github.com/dholab/cyclospora-in-wastewater-metagenomics/blob/main/01-identify-cyclospora-specific-kmers/results/cyclospora_cayetanensis_rrna_kmer_manifest.tsv).

The 5,561 candidates were then passed through three successive filters, (1) removing 31-mers present in the ribosomal RNA of other organisms, (2) 31-mers present in the ribosomal RNA of other *Cyclospora* species, and (3) 31-mers too repetitive to be informative. To remove 31-mers in rRNA from other organisms, we downloaded SILVA 138.2 SSURef_NR99 and LSURef_NR99, which contain 18S and 28S sequence from across the tree of life with near-duplicates removed at 99% identity, together with Rfam 15.1 families RF00001 and RF00002, which include the 5S and 5.8S that SILVA does not contain. We excluded the 9 *C. cayetanensis* rRNA sequences from SILVA and these nine plus [329 C. cayetanensis rRNA accessions](https://github.com/dholab/cyclospora-in-wastewater-metagenomics/blob/main/01-identify-cyclospora-specific-kmers/config/target_rrna_accessions.txt) from Rfam (src/rrna_bait/sources.py). The k-mers in the non-*C. cayetanensis* rRNAs were then enumerated with [meryl count k=31](https://github.com/dholab/cyclospora-in-wastewater-metagenomics/blob/main/01-identify-cyclospora-specific-kmers/scripts/build_rrna_bait.sh#L408-L413) as described above. We then removed every *C. cayetanensis* 31-mer that also appears in SILVA or Rfam using [meryl union](https://github.com/dholab/cyclospora-in-wastewater-metagenomics/blob/main/01-identify-cyclospora-specific-kmers/scripts/build_rrna_bait.sh#L414-L417) after merging the SILVA and Rfam 31-mer sets into a single “background 31-mer set”. [meryl difference](https://github.com/dholab/cyclospora-in-wastewater-metagenomics/blob/main/01-identify-cyclospora-specific-kmers/scripts/build_rrna_bait.sh#L418-L421) subtracts this background 31-mer set from the *C. cayetanensis* 31-mers. This left 1,839 k-mers that occur in *C. cayetanensis* ribosomal RNA and in no ribosomal RNA sequences from SILVA or Rfam.

Next we removed 31-mers shared between *C. cayetanensis* and the other species of its own genus. The rRNA of those relatives are largely missing from SILVA because the SSU Ref only considers sequences 1,200 bases or longer, and NR99 then dereplicates what remains at 99% identity, which is finer than the differences between congeneric *Cyclospora*. There are [94 additional Cyclospora rRNA sequences we retrieved](https://github.com/dholab/cyclospora-in-wastewater-metagenomics/blob/main/01-identify-cyclospora-specific-kmers/config/other_cyclospora_accessions.txt) from NCBI Genbank, mostly partial amplicons deposited from genotyping and barcoding studies. We counted their 31-mers with [meryl count k=31](https://github.com/dholab/cyclospora-in-wastewater-metagenomics/blob/main/01-identify-cyclospora-specific-kmers/scripts/build_rrna_bait.sh#L412-L413), added that set to the SILVA and Rfam background with [meryl union](https://github.com/dholab/cyclospora-in-wastewater-metagenomics/blob/main/01-identify-cyclospora-specific-kmers/scripts/build_rrna_bait.sh#L422-L426), and subtracted the combined background from the target with [meryl difference](https://github.com/dholab/cyclospora-in-wastewater-metagenomics/blob/main/01-identify-cyclospora-specific-kmers/scripts/build_rrna_bait.sh#L427-L430) a second time. That removed 169 k-mers and left 1,670.

For the third filter, we applied [Deacon](https://github.com/bede/deacon), using [deacon index build -k 31 -w 1 -e 0.6](https://github.com/dholab/cyclospora-in-wastewater-metagenomics/blob/main/01-identify-cyclospora-specific-kmers/scripts/build_rrna_bait.sh#L477-L481), which scores each k-mer by the Shannon entropy of its base composition, scaled so that 1.0 is four equally frequent bases and 0 is a homopolymer, and drops any k-mer scoring below 0.6. Deacon retained all 1,670 k-mers.

The surviving baits are distributed unevenly across the four genes. 28S rRNA, the longest of the four loci, supplied 1,550 of the 1,670 31-mers. 45% of the 28S candidates survived, compared with 5% of the 18S candidates. This may mean that 28S is the better target for a diagnostic assay, but it may also reflect uneven coverage of the background: SILVA’s large-subunit reference set is roughly a third the size of its small-subunit set, 70.6 MB against 201.1 MB compressed (config/sources.tsv), so a 28S candidate had fewer opportunities to be removed.

### 2.3 Exact taxonomic screening retained 1,464 baits

An apparently *C. cayetanensis*-specific 31-mer absent from both SILVA and Rfam could still occur in an unannotated region of a genome, in a metagenomic assembly, or in an organism whose rRNA has never been cataloged in SILVA/Rfam (e.g., rRNA sequences shorter than 1,200 bases). We therefore screened all 1,670 baits against the 28 July 2025 NCBI `core_nt` release with [blastn -word_size 31 -ungapped -perc_identity 100 -qcov_hsp_perc 100 -dust no](https://github.com/dholab/cyclospora-in-wastewater-metagenomics/blob/main/REPRODUCING.md#repeating-the-core-nt-search), which reports only exact, full-length matches to a 31-mer.

A bait was rejected if any exact 31-of-31 match belonged to a taxon other than *C. cayetanensis* (taxid 88456), or matched a sequence carrying no taxon assignment at all (src/rrna_bait/core_nt.py). The screen rejected 206 baits and retained 1,464, as recorded in results/cyclospora_cayetanensis_core_nt_validation.tsv. The retained set contains 61 baits from 18S, 1,372 from 28S, 31 from 5S, and none from 5.8S (cyclospora_cayetanensis_rrna_core_nt_validated_baits.fasta).

Mapping the retained baits back to the seven target sequences revealed strong spatial clustering (cyclospora_cayetanensis_rrna_kmer_manifest.tsv). The single 28S target contains 29 runs of baits beginning at consecutive positions, the longest containing 301 baits across 331 nucleotides. Each 18S target contains three runs. One 5S target contains a 31-bait run across 61 nucleotides; the other two 5S targets contain none of the retained baits.

We then built a Deacon filtering index from these 1,464 31-mers using [deacon index build -k 31 -w 1 -e 0](https://github.com/dholab/cyclospora-in-wastewater-metagenomics/blob/main/01-identify-cyclospora-specific-kmers/scripts/finalize_core_nt_validation.sh#L96-L99). Setting `w=1` makes every 31-mer its own minimizer, so nothing is subsampled and all diagnostic 31-mers are in the index.

### 2.4 Setting a calibration threshold of 24 diagnostic 31-mers before a read counts as *Cyclospora*

To determine how many 31-mers a read must carry before it can be confidently classified as *Cyclospora*, we used 293 public wastewater metagenomic SRA runs from BioProject PRJNA1247874, collected from 27 sewersheds between June 1 and August 31, 2025. We screened the read sets with the 1,464-bait index at `deacon filter -a 1 -r 0`. Of the 293 runs, 232 returned at least one bait-bearing read, yielding 108,474 reads carrying at least one bait. Each distinct read sequence was submitted in full to BLASTN against the same `core_nt` database used to validate the baits. BLASTN performs local alignment, and we imposed no minimum query-coverage criterion. We classified each read by comparing its highest local-alignment bit score to a taxon in the declared 26-taxid *Cyclospora* scope with its highest score outside that scope. Scores differing by less than 0.1 bits were retained as ties.

The presence of a single 31-mer is not sufficiently specific on its own. A read can carry a bait for which the screened `core_nt` release contains no exact non-target match and still produce a stronger local alignment to another organism. At a threshold of one bait, the calibration contained 1,284 target reads, 107,040 non-target reads, 123 ties, and 27 reads with no `core_nt` hit.

Non-target or tied reads remained through a bait count of 23 (Table 3). At a minimum count of 24, 853 target-classified reads and no non-target, tied, or no-hit reads remained. We therefore used an absolute Deacon threshold of 24 for our scan across all public samples in SRA BioProject PRJNA1247874.

**Table 1:**
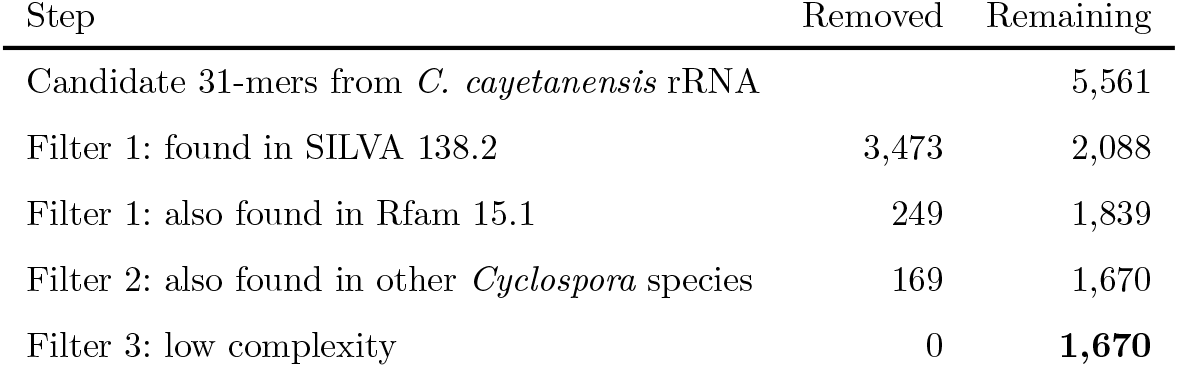
Filtering removes non-specific 31-mers. Each row counts only the k-mers that the rows above it had not already removed, so no k-mer is counted twice. The complete decisions are recorded in the k-mer manifest.

**Table 2:** The retained bait set is dominated by 28S rRNA. Percentages are relative to the 1,464 baits that passed taxonomic exact-match screening.

| rRNA | Retained baits | Share of final set |
| --- | --- | --- |
| 18S | 61 | 4.2% |
| 28S | 1,372 | 93.7% |
| 5S | 31 | 2.1% |
| 5.8S | 0 | 0% |

**Table 3:**
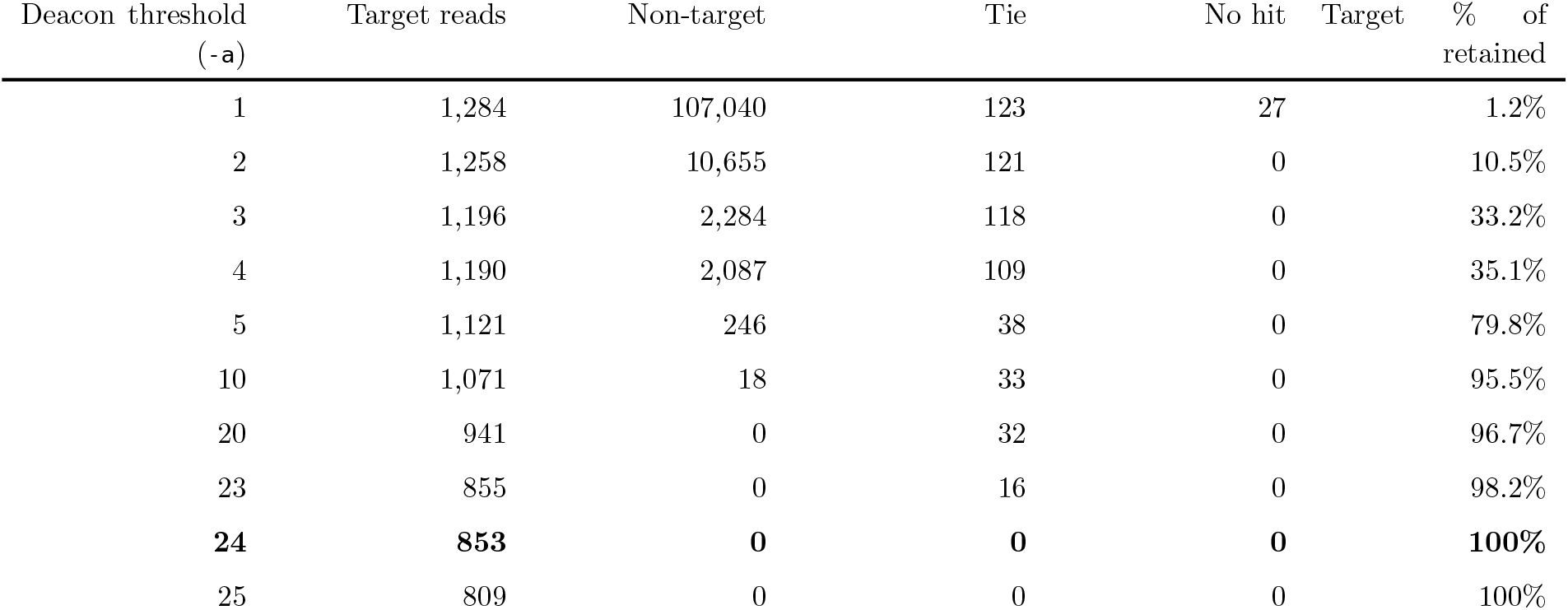
Specificity for Cyclospora improves as the threshold for diagnostic 31-mers per-read rises. Each row counts the calibration reads carrying at least that many diagnostic 31-mers, classified by comparing the best local-alignment bit scores within and outside the declared *Cyclospora* target scope. Selected thresholds from `threshold_read_counts.tsv`, which lists every value from 1 to 121.

Mapping the selected threshold back to the target sequences showed that 16 of the 29 contiguous 28S runs contain at least 24 baits (bait_calibration_summary.json). Of the 3,338 possible 150-nucleotide windows in the 28S target, 2,061 (61.7%) contain at least 24 baits and thus meet the threshold. Each 18S target contains one threshold-capable run, and approximately 4.6% of its possible 150-nucleotide windows meet the threshold. The 31-bait 5S run can also satisfy the threshold.

Of the 108,474 read observations, 96,440 (88.9%) carried one bait. Non-target reads had a median of one bait and a maximum of 15, whereas target-classified reads had a broad distribution with a median of 42 and a range of 1–121. Bait utilization was correspondingly heavy-tailed before thresholding: 138 baits were unused, 23 appeared in one read observation, and the ten most frequently observed baits accounted for 65.7% of all bait–read incidences. The most frequent was an 18S bait with no exact `core_nt` hit, which occurred in 95,196 non-target observations and no target observations. Those reads were predominantly classified by their remaining sequence as uncultured fungal 18S. Among the 853 target observations passing threshold 24, utilization was much more even: the ten most frequent baits contributed 2.5% of incidences.

The target-supporting local alignments were also high identity. Across the 853 threshold-passing observations, the median identity of the highest-scoring target alignment was 100%, 87.9% were at least 99% identical, and the median query coverage was 99%; 72.0% covered at least 95% of the read. These observations represented 183 unique read sequences. At the exact boundary of 24 baits, 44 observations collapsed to three unique sequences whose target alignments were 98.6–99.2% identical over 81–97% of the read and exceeded the best non-target score by 36– 67 bits.

Of the 1,464 retained baits, 130 had no exact match of any kind in `core_nt`. One is the 18S bait described above. The other 129 are 28S baits spanning four sites where the 28S target differs by one to three bases from the *C. cayetanensis* 28S records in `core_nt`, so every 31-mer crossing a difference lost its exact match. Removing all 130 left the selected threshold at 24 and retained 852 of the 853 target observations.

### 2.5 *Cyclospora* recurs seasonally across the public wastewater record

We screened 2,333 public [CASPER](https://securebio.org/casper/) wastewater metagenomes in NCBI SRA BioProject PRJNA1247874 from the [frozen accession roster](https://github.com/dholab/cyclospora-in-wastewater-metagenomics/blob/main/02-screen-wastewater-metagenomes/config/wastewater_cohort.tsv). The samples were collected between 2023-12-26 and 2026-06-30 and screened using the calibrated Deacon filter (`deacon filter -a 24 -r 0`) against the 1,464-bait index. After recounting each emitted read separately, 81 runs (3.5% of all screened runs) contained putative *C. cayetanensis* reads. To focus our analysis on sewersheds with longitudinal data, we restricted the analysis to the 30 sewersheds sampled at 10 or more timepoints (2,328 runs); 24 of those 30 sewersheds contained at least one positive run (Figure 1)

As expected from the seasonal epidemiology of cyclosporiasis Casillas et al., 2019, the detections in wastewater are strongly seasonal. The public record spans two cyclosporiasis seasons and *C. cayetanensis* rises in both: detection climbs from near zero in spring to roughly one run in nine across June, July and August of 2025, falls back to near zero through the winter, and rises again in the early summer of 2026. That the same seasonal pattern appears in two independent summers is strong evidence that these detections track community infection rather than sequencing artifacts.

**Figure 1:**
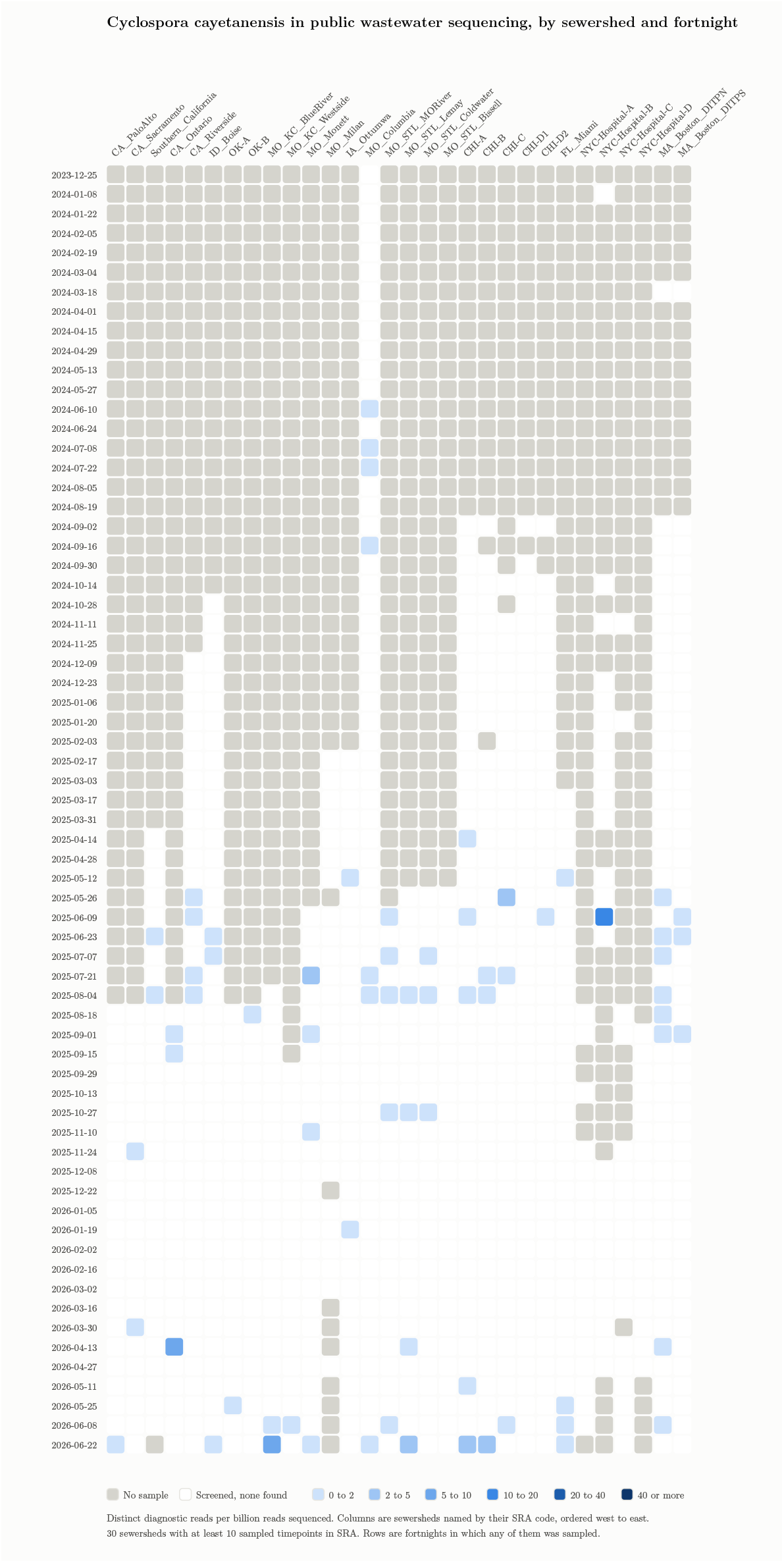
*Cyclospora cayetanensis* in public wastewater sequencing data, by sewershed and fortnight. In the [interactive version](https://dholab.github.io/cyclospora-in-wastewater-metagenomics/#figure-1), hovering a cell reports its distinct read count, sequencing depth, and contributing runs, and clicking a cell with detections opens those same distinct reads as FASTA, to read, copy, or download. Color shows distinct *C. cayetanensis* reads per billion reads sequenced, where distinct is the count of unique read sequences per run, so PCR and optical copies are collapsed; the denominator is the sum of input reads across the runs contributing to each cell. Each column is a separate sewershed, geographically ordered west to east. Gray marks a fortnight with no sample and white a run screened with no *C. cayetanensis* reads found.

## 3 Discussion

Here we demonstrate that wastewater metagenomic sequencing can identify samples with rRNA reads that are more similar to *C. cayetanensis* than any other taxa in NCBI core-nt. The most parsimonious explanation is that these reads are, in fact, from *C. cayetanensis* parasites shed into wastewater. However, we cannot formally exclude the possibility that some of these detections are associated with as yet undiscovered *Cyclospora* species that are not represented in core-nt or other databases. Moreover, it is impossible to differentiate locally acquired from travel-associated cases based on wastewater data alone, and travel-associated cases may be less concerning for public health. Our intentionally conservative threshold for assigning reads to *C. cayetanensis* may also result in false negatives, especially when levels of *C. cayetanensis* are low. Nevertheless, it is reasonable to cautiously assert that this approach may be useful for monitoring Cyclospora outbreaks and could provide situational awareness to clinicians and other stakeholders when *C. cayetanensis* is detectable in their communities.

The 1,464-bait index is effectively a 28S assay, and much of its detection capacity occurs in contiguous runs of overlapping 31-mers. This clustering likely reflects regions of rRNA that are conserved within *C. cayetanensis* but diverge from the sequences represented in the background databases. We have not evaluated whether similarly concentrated diagnostic runs emerge for other taxa. The threshold of 24 should therefore be understood as a minimum amount of matching sequence rather than 24 independent loci: 24 consecutive 31-mers can fit within a 54-nucleotide tract. A substitution near the center of such a tract can remove all 24 matches, creating a potential false negative, while an unrepresented organism sharing the tract can satisfy the threshold. The concentration of baits in 28S may likewise reflect genuine locus-specific divergence, uneven reference coverage, or both. We would thus not take this as evidence that 28S is intrinsically more diagnostic than other regions. Rather, 28S happened to be the locus offering most diagnostic k-mers given the sequences represented in the background we screened against.

Indeed, any reference will suffer from representation gaps. As an example, one of our baits that lacked an exact match in Core NT occurred in more than 95,000 reads whose strongest local alignments supported other organisms. `core_nt` representation is uneven across taxa, and using the same release for bait screening and read classification means that the two stages share its blind spots. Specificity should therefore be interpreted relative to the 28 July 2025 release and the organisms observed in the calibration panel, not as universal absence from nature. Reassuringly, a requirement of 24 baits buffered the analysis against off-target baits, and the threshold was unchanged when we removed the 130 baits lacking any exact match in Core NT. This empirical calibration is more informative than treating a negative database search as proof of specificity, but related coccidia missing from the reference collection remain the most plausible source of false-positive attribution.

In the summer of 2026 the United States experienced a cyclosporiasis season large enough to command national attention. By the middle of July the general medical press was describing an outbreak of “explosive diarrhoea” sweeping the country (Brown, 2026), followed within days by coverage asking whether people should avoid fruit and vegetables (Wise, 2026) and by a clinical primer for patients in *JAMA* (Linder and Malani, 2026). A nationwide [CBS News survey](https://www.cbsnews.com/news/cyclospora-outbreak-fda-opinion-poll) conducted from July 22-24 found that approximately 40% of Americans are buying or eating less produce in response to this outbreak. We have analyzed more recent wastewater metagenomic data from the summer of 2026 (i.e., data not yet in SRA) that we are not including in this analysis. This data has been shared with public health stakeholders in the jurisdictions where we collect wastewater for the CASPER project. As of this writing we are not including Cyclospora detection data on our public dashboard wastewater metagenomic dashboard, consistent with our approach to other high-consequence pathogens (e.g, Measles virus, MPXV).

The approach we describe here could be a useful blueprint for early detection of *C. cayetanensis* and focusing messaging specifically on at-risk populations, in part because conventional clinical surveillance for this parasite is slow. The median interval from a patient’s illness onset to their diagnosis is 20 days, and the median interval from that diagnosis to CDC notification can take an additional month (Casillas et al., 2019; PMID 31002104). This means a typical case in the national count lags symptoms by weeks, if not a month or more. Moreover, a case is counted only if a clinician thinks to order a *Cyclospora* test, which is not part of a routine examination for ova and parasites and appears in only one of the several commercial multiplex gastrointestinal panels (Casillas et al., 2019). Moreover, clinical case ascertainment requires infected people to seek care, which in turn requires access to medical care and symptoms severe enough to prompt a visit. [Behavior-independent wastewater testing](https://www.amazon.com/Everyone-Poops-Taro-Gomi/dp/1797202642) does not depend on either. The same logic has already played out for measles: in October 2025 the Hawaii Department of Health reported genotype D8 measles virus in a wastewater sample from East Kauai at a time when no suspected measles case had been identified on the island (Hawaii News Now, 22 October 2025).

Moreover, the selective “spearfishing” approach to *C. cayetanensis* detection in wastewater metagenomic datasets could be applied to other pathogens and their hosts. Sample preparation for these datasets are optimized for virus recovery, and a serendipitous benefit is that most samples carry a substantial mature rRNA fraction. To date, most of our effort has focused on virus detection and non-viral microbial detection with tools like GOTTCHA2, whose signature-based approach is described in Freitas et al., 2015, and which rely on subsampled k-mer reference databases that are less sensitive for rRNA detections than the exhaustive k-mers used in this analysis. Indeed, GOTTCHA2 did identify *C. cayetanensis* reads from some of these samples, albeit less sensitively. It is trivial to generate 31-mers for pathogens of interest to determine if there are enough useful k-mers for classification. In some cases, the answer will be ‘no.’ For example, applying the same approach to *Bordetella pertussis* and *Bordetella parapertussis* did not yield enough diagnostic 31-mers to be useful (unpublished observation). However, this approach can be applied on a case-by-case basis with other targets where it might be fruitful. An advantage of the longitudinal wastewater data, collected from the same communities over multiple years, is that it can be mined to address questions like this that are only obvious with the benefit of hindsight. It also underscores the value of expanding the locations and number of sites where data is being obtained to improve the resolution of how pathogens move through time and geographic space.

## 4 Methods

### 4.1 Bait design and taxonomic validation

We aligned [nine Cyclospora cayetanensis rRNA records](https://github.com/dholab/cyclospora-in-wastewater-metagenomics/blob/main/01-identify-cyclospora-specific-kmers/config/target_queries.tsv)—one 18S, one 28S, one 5.8S, and six 5S records—to two pinned RefSeq assemblies, GCF_002999335.1 (CcayRef3) and GCF_000769155.1 (ASM76915v2), using BLAST+ 2.16.0 (`blastn -task megablast`). We retained alignments with at least 98% identity and 90% query coverage. After extracting the matching genomic intervals and deduplicating sequences, we obtained seven putative mature rRNAs: two 18S, one 28S, three 5S, and one 5.8S.

Using [meryl 1.4.2](https://github.com/marbl/meryl), we enumerated all 31-mers in these sequences and their reverse complements and retained one copy of each. We discarded any found in SILVA 138.2 SSURef_NR99 or LSURef_NR99, Rfam 15.1 families RF00001 or RF00002, or [94 other Cyclospora rRNA records retrieved from GenBank](https://github.com/dholab/cyclospora-in-wastewater-metagenomics/blob/main/01-identify-cyclospora-specific-kmers/config/other_cyclospora_accessions.txt). We then used [Deacon 0.15.0](https://doi.org/10.1101/2025.06.09.658732) to remove low-complexity 31-mers, scoring each by the Shannon entropy of its base composition scaled to the range 0 to 1 and dropping those below 0.6; none were. The remaining 1,670 baits were searched for exact 31-of-31 matches against the 28 July 2025 `core_nt` release. We removed any bait with an exact match assigned outside *C. cayetanensis* taxid 88456 or lacking a taxonomic assignment. The final 1,464 baits were compiled into a Deacon index with `k=31`, `w=1`, and `e=0`. Software and reference versions are recorded in the [locked analysis environment](https://github.com/dholab/cyclospora-in-wastewater-metagenomics/blob/main/01-identify-cyclospora-specific-kmers/pixi.lock) and [source manifest](https://github.com/dholab/cyclospora-in-wastewater-metagenomics/blob/main/01-identify-cyclospora-specific-kmers/config/sources.tsv).

### 4.2 Detection-threshold calibration

We calibrated a minimum-k-mer detection threshold for *Cyclospora cayetanensis* using 293 wastewater metagenomic SRA runs collected during summer 2025. We first screened with Deacon at a threshold of one bait. Because Deacon evaluates paired reads jointly, we recounted the baits in each read before choosing the threshold. Within each run, we collapsed exact duplicate fragments, defining duplicates as read pairs for which both mates had the same sequence or reverse complement. We then discarded any mate carrying no bait. To avoid redundant database searches, each unique retained read sequence was queried once with BLASTN against the same `core_nt` release, and its classification was mapped back to the corresponding reads. We classified reads by comparing the highest local-alignment bit score among matches assigned to a [predefined target set of 26 Cyclospora taxids](https://github.com/dholab/cyclospora-in-wastewater-metagenomics/blob/main/02-screen-wastewater-metagenomes/results/calibration/cyclospora_genus_taxids.txt) with the highest score among all other matches. Differences below 0.1 bits were treated as ties, and no minimum query coverage was required. We selected the lowest threshold retaining at least one target-classified read and no non-target, tied, or no-hit reads: 24 baits per read.

### 4.3 BioProject screening and longitudinal visualization

We analyzed public CASPER metagenomes from BioProject PRJNA1247874, generated from filtered wastewater concentrated by polyethylene glycol precipitation as described by Rushford et al. (2025) and Justen et al. (2026). We supplied the [frozen SRA accession roster](https://github.com/dholab/cyclospora-in-wastewater-metagenomics/blob/main/02-screen-wastewater-metagenomes/config/wastewater_cohort.tsv) to NVD 3.5.0 (revision `bd421147bba2a8ba22aafffb5dbf61c47d241feb`, clean). NVD streamed each run from SRA and screened it with Deacon 0.16.0 using the validated index, an absolute threshold of 24, and a relative threshold of zero. Although Deacon evaluates paired reads jointly, downstream analysis applied the threshold separately to every emitted read. Within each run, identical complete read sequences were counted once, with reverse complements treated as identical. We grouped runs by sewershed in 14-day intervals beginning on Mondays, summed the number of

retained reads and input reads, and reported the ratio as reads per billion input reads. Figure 1 includes surveillance sites sampled on at least ten dates and orders them by longitude from west to east.

## 5 Data availability

All sequencing analyzed here is public in NCBI SRA under BioProject PRJNA1247874

| What | Where |
| --- | --- |
| Source sequencing | NCBI SRA BioProject PRJNA1247874 |
| Per-run results, all 2,333 screened SRA runs | sra_sample_summary.tsv |
| Diagnostic reads, one FASTA member per positive run | diagnostic_reads.tar.gz |
| The sewershed codes, with coordinates and role | casper_sites.tsv |
| The matrix plotted in Figure 1 | site_fortnight_matrix_per_billion.tsv |
| Threshold calibration evidence for 108,474 reads carrying at least one bait | 02-screen-wastewater-metagenomes/results/calibration/ |
| The bait sets | 01-identify-cyclospora-specific-kmers/baits/ |
| Bait design and core-nt validation evidence | 01-identify-cyclospora-specific-kmers/results/ |

## 6 Acknowledgements

This work was performed as part of the [Lungfish Research](https://lung.fish/) project funded by [Inkfish](https://ink.fish/).

The wastewater metagenomics datasets are generated by the CAPSER collaboratory. We would like to thank all of the participating sewersheds who contribute wastewater samples weekly and who make this sort of analysis possible.

Computing was performed on the [UW-Madison Center for High Throughput Computing](https://chtc.cs.wisc.edu/) cluster. Claude Code, Claude Opus v5 and v4.8, and ChatGPT v5.6 Sol were used for research, data analysis, and text drafting.

## References

Ambrose, K., Hamilton-Seth, R., Jackson, M., Sigler, R., Fels Elliott, D.R. (2025) Chronic Cyclospora infection in a heart transplant patient with intestinal malabsorption: a case report. ASM Case Rep 1(6). doi:10.1128/asmcr.00062-25 · PMID 41244276 · PMC12584178

Ashford, R.W. (1979) Occurrence of an undescribed coccidian in man in Papua New Guinea. Ann Trop Med Parasitol 73(5):497–500. doi:10.1080/00034983.1979.11687291 · PMID 534451

Banasik, E., Dobrowolska, A., Woźnicka-Leśkiewicz, L., Eder, P. (2025) Diagnostic challenges of cyclosporiasis in chronic diarrhea: a case study. Microorganisms 13(9):2209. doi:10.3390/microorganisms13092209 · PMID 41011540 · PMC12472525

Barratt, J.L.N., Shen, J., Houghton, K., Richins, T., Sapp, S.G.H., Cama, V., Arrowood, M.J., Straily, A., Qvarnstrom, Y. (2023) Cyclospora cayetanensis comprises at least 3 species that cause human cyclosporiasis. Parasitology 150(3):269–285. doi:10.1017/S003118202200172X · PMID 36560856 · PMC10090632

Brown, C. (2026) Cyclosporiasis: as “explosive diarrhoea” sweeps US, what’s behind the outbreak and what is Trump’s role? BMJ 394:e100317. doi:10.1136/bmj-2026-100317 · PMID 42468983

Casillas, S.M., Hall, R.L., Herwaldt, B.L. (2019) Cyclosporiasis surveillance — United States, 2011–2015. MMWR Surveill Summ 68(3):1–16. doi:10.15585/mmwr.ss6803a1 · PMID 31002104 · PMC6476303

Constantinides, B., Lees, J.A., Crook, D.W. (2025) Deacon: fast sequence filtering and contaminant depletion. bioRxiv 2025.06.09.658732. doi:10.1101/2025.06.09.658732 · preprint, not peer reviewed

Durigan, M., Murphy, H.R., da Silva, A.J. (2020) Dead-end ultrafiltration and DNA-based methods for detection of Cyclospora cayetanensis in agricultural water. Appl Environ Microbiol 86(23):e01595–20. doi:10.1128/AEM.01595-20 · PMID 32948525 · PMC7657621. Published as BAM Chapter 19c by the FDA.

Freitas, T.A.K., Li, P.-E., Scholz, M.B., Chain, P.S.G. (2015) Accurate read-based metagenome characterization using a hierarchical suite of unique signatures. Nucleic Acids Res 43(10):e69. doi:10.1093/nar/gkv180 · PMID 25765641 · PMC4446416

Goetzman, J., Carter, A., Oliveira, A., Ingram, L.A. (2025) Outbreak of cyclosporiasis among patrons of a Mexican-style restaurant — Limestone County, Alabama, May–June 2023. MMWR Morb Mortal Wkly Rep 74(13):217–221. doi:10.15585/mmwr.mm7413a1 · PMID 40244909 · PMC12005484

Hall, N.B., Chancey, R.J., Keaton, A.A., Heines, V., Cantu, V., Vakil, V., Long, S., Short, K., Franciscus, E., Wahab, N., Gieraltowski, L., Straily, A. (2020) Cyclosporiasis epidemiologically linked to consumption of green onions: Houston metropolitan area, August 2017. J Food Prot 83(2):326–330. doi:10.4315/0362-028X.JFP-19-254 · PMID 31961230 · PMC10130782

Hofstetter, J., Arfken, A., Kahler, A., Qvarnstrom, Y., Rodrigues, C., Mattioli, M. (2024) Evaluation of coccidia DNA in irrigation pond water and wastewater sludge associated with Cyclospora cayetanensis 18S rRNA gene qPCR detections. Microbiol Spectr 12(8):e0090624. doi:10.1128/spectrum.00906-24 · PMID 38916361 · PMC11302338

Justen, L.J., Rushford, C., Hershey, O.S., Floyd-O’Sullivan, R., Grimm, S.L., Bradshaw, W.J., Bhasin, H., Rice, D.P., Stansifer, K., Faraguna, J.D., McLaren, M.R., Zulli, A., Tovar-Mendez, A., Copen, E., Shelton, K.K., Amirali, A., Kannoly, S., Pesantez, S., Stanciu, A., Quiroga, I.C., Silvera, L., Greenwood, N., Bongiovi, B., Walkins, A., Love, R., Lening, S., Patterson, K., Johnston, T., Hernandez, S., Benitez, A., McCarley, B.J., Engelage, S., Pillay, S., Calender, C., Herring, B., Robinson, C. (2026) Deep untargeted wastewater metagenomic sequencing from sewersheds across the United States. medRxiv, version 2 posted 18 March 2026. doi:10.64898/2026.03.05.26345726 · preprint, not peer reviewed

Kahler, A.M., Hofstetter, J., Arrowood, M., Peterson, A., Jacobson, D., Barratt, J., da Silva, A.L.B.R., Rodrigues, C., Mattioli, M.C. (2024) Sources and prevalence of Cyclospora cayetanensis in southeastern U.S. growing environments. J Food Prot 87(7):100309. doi:10.1016/j.jfp.2024.100309 · PMID 38815808 · PMC11288253

Kitajima, M., Haramoto, E., Iker, B.C., Gerba, C.P. (2014) Occurrence of Cryptosporidium, Giardia, and Cyclospora in influent and effluent water at wastewater treatment plants in Arizona. Sci Total Environ 484:129–136. doi:10.1016/j.scitotenv.2014.03.036 · PMID 24695096

Linder, K.A., Malani, P.N. (2026) What is cyclosporiasis? JAMA, published online 21 July 2026. doi:10.1001/jama.2026.14866 · PMID 42479478

McCaughan, K.J., Kniel, K.E. (2026) Current knowledge and future directions for Cyclospora cayetanensis research and its surrogates. Compr Rev Food Sci Food Saf 25(1):e70327. doi:10.1111/1541-4337.70327 · PMID 41347294

Ortega, Y.R., Sterling, C.R., Gilman, R.H., Cama, V.A., Díaz, F. (1993) Cyclospora species — a new protozoan pathogen of humans. N Engl J Med 328(18):1308–1312. doi:10.1056/NEJM199305063281804 · PMID 8469253

Ortega, Y.R., Gilman, R.H., Sterling, C.R. (1994) A new coccidian parasite (Apicomplexa: Eimeriidae) from humans. J Parasitol 80(4):625–629. PMID 8064531 (no DOI registered)

Rushford, C., Gregory, D.A., Copen, E.E., Naydenov, J., Tovar-Mendez, A., Li, P.-E., Kantor, R.S., Shakya, M., Ruth, N., Chain, P.S.G., Hsieh, H.-Y., Hunter, T., Kome, J., Frank, M., Lyddon, T.D., Kaufman, J., O’Connor, D.H., Johnson, M.C. (2025) Untargeted longitudinal ultra deep metagenomic sequencing of wastewater provides a comprehensive readout of expected and unexpected viral pathogens. medRxiv, version 2 posted 27 January 2026. doi:10.1101/2025.10.27.25338874 · preprint, not peer reviewed

Shen, J., Cama, V.A., Jacobson, D., Barratt, J., Straily, A. (2025) Cyclospora genotypic variations and associated epidemiologic characteristics, United States, 2018–2021. Emerg Infect Dis 31(2):256–266. doi:10.3201/eid3102.240399 · PMID 39983699 · PMC11845137

Sturbaum, G.D., Ortega, Y.R., Gilman, R.H., Sterling, C.R., Cabrera, L., Klein, D.A. (1998) Detection of Cyclospora cayetanensis in wastewater. Appl Environ Microbiol 64(6):2284–2286. doi:10.1128/AEM.64.6.2284-2286.1998 · PMID 9603852 · PMC106316

Wise, J. (2026) Cyclosporiasis: should people avoid fruit and vegetables? BMJ 394:e100324. doi:10.1136/bmj-2026-100324 · PMID 42476611

